# ColMA/PEGDA Bioink for Digital Light Processing 3D Printing in Biomedical Engineering

**DOI:** 10.1101/2023.01.08.523134

**Authors:** Jishizhan Chen

## Abstract

**Introduction:** Digital light processing (DLP) represents a rapid approach to constructing 3D structures with superior resolution. However, it imposes demanding requirements on the properties of bioink. Gelatine methacryloyl has long been the major option but results in limited mechanical properties. The development of collagen-based bioink provides a wider stiffness adjustment range, native bioactivities, and versatility in biomedical engineering applications.

**Method:** Collagen solution was obtained via enzymolysis and ultrafiltration and then subjected to methacrylation. The photocrosslinkable bioink comprises collagen methacryloyl (ColMA), poly(ethylene glycol) diacrylate (PEGDA), acetic acid, yellow food dye, and photoinitiator LAP. The 3D structures were fabricated utilising a commercial DLP printer with 405 nm visible light.

**Results:** Purified type I collagen can be rapidly obtained via the developed process, and methacrylation is optimised for collagen with much less addition of methacrylic anhydride (MAA) and a high degree of substitution. The ColMA/PEGDA bioink is translucent and low viscosity and is suitable for DLP 3D printing. The printed scaffolds reached a compressive modulus over 100 kPa with 0.6 wt% collagen. Sharp-edged and fine structures (∼500 μm) were obtained by printing. The hydrogels show tunable mechanical properties by adjusting the concentration of the ColMA component. A series of models were fabricated to test the printability, including ear, cube with channels, and scaffolds, which display porous structures with pore sizes of 50 – 150 μm.

**Conclusions:** An optimised collagen-based bioink fabrication protocol was proposed for the DLP technique, covering steps from collagen extraction to ColMA/PEGDA bioink formulation and printing. Bioink with tunable mechanical properties is suitable for DLP printing. High-resolution structures can potentially be utilised for various biomedical engineering applications.

## 1. Introduction

3D printing, or additive manufacturing, is a technique that can print out and solidify biological materials into any geometric structure[1]. The patient-customised capability turns it into a popular approach in biomedical engineering. According to the Web of Science[2], the number of annual publications containing the key word ‘3D printing’ has grown from 314 to 10,574 (Supplementary Figure 1) in the past decade. The main 3D printing technologies include extrusion-based[3], inkjet-based[4], and laser-assisted[5] printing. Extrusion-based printing relies on mechanical or pneumatic power to squeeze biological ink through the nozzle. The printing resolution is significantly limited by the diameter of the nozzle[6]. Structural collapse, especially when printing large or high structures, is another obvious hurdle using collagen or gelatin-based bioink due to insufficient viscosity[7]. Inkjet-based printing propels bioink through heating or piezoelectricity at a certain frequency to form a structure. It requires bioink with low viscosity that can be dispensed onto the platform and then quickly solidified. However, the triple helix structure of collagen is sensitive to temperature and may denature with increasing temperature, leading to reduced bioactivity and uncontrollable changes in viscosity. Additionally, in the case of low viscosity ink mixed with cells, the cells tend to settle to the bottom of the ink cartridge due to gravity in static conditions[8]. The cells may be dispersed unevenly in the printed structure or easily become clogged within the nozzle. Hence, inkjet-based printing is not ideal for collagen-based bioink. Laser-assisted printing uses a laser to replace heating or piezoelectricity in inkjet-based printers. The bioink is cured by laser, which usually has a short wavelength and takes a relatively long time. The laser damages DNA and generates much heat, causing harm to cells in the bioink[9].

Apart from the above-described conventional 3D printing technologies, novel digital light processing (DLP) [10] appears to be an optimal technique for sensitive bioink. The projector is the core unit of a DLP printer. By projecting 2D images into the light-sensitive liquid sample pool, curing is processed in a layer-by-layer manner. DLP can be performed at low temperature using visible light. Moreover, the printing resolution is no longer limited by the size of the nozzle but by the pixels of the image, which allows higher resolution and faster speed than nozzle-based printing[11]. These characteristics ensure the maximal preservation of material bioactivity and cell viability. Despite these advantages, DLP printing is performed on an upside-down printing platform, which highly relies on the surface tension of the liquid bioink. This feature raises rigorous requirements for the viscosity of the bioink.

Collagen and its thermally denatured product gelatin have been widely utilised in 3D printing because they provide similar components to human tissues such as bones, skin, blood vessels, ligaments, and cartilage. To date, the bioink specialised for DLP printing is still dominated by gelatin methacryloyl (GelMA). This is mainly attributed to the simpler fabrication process and printing conditions of gelatin. However, the low mechanical properties of gelatin-based DLP bioink limit its application. The collagen-based bioink encounters obstacles such as a long production cycle, not being optimised for DLP printing, and uncontrollable crosslinking depth. Some obviously contradictory reaction conditions further impede the fabrication of collagen-based DLP bioink. To overcome these challenges, we developed a protocol for type I collagen methacryloyl (ColMA)/poly(ethylene glycol) diacrylate (PEGDA) bioink specified for DLP. It shows characteristics of rapid bioink manufacture and DLP suitability. A series of structures were fabricated to test the printability.

## 2. Materials and methods

Bovine tendon was purchased from the local market. Pepsin from porcine gastric mucosa (lyophilized powder, ≥3200 units/mg protein), Ultra Centrifugal Filter Units (100 kDa cut-off filter), methacrylic anhydride (MAA), and Collagen Assay Kit were purchased from Sigma-Aldrich (Missouri, US). The XCell SureLock™ Mini-Cell Electroblotting Unit, Rapid Gold BCA Protein Assay Kit, and all the other reagents and consumables were purchased from Thermo Fisher Scientific (Massachusetts, USA) unless otherwise specified.

### 2.1 Extraction and purification of type I collagen

All solutions were equilibrated to 4°C before use, and all operations were conducted at 4°C unless otherwise specified. Soft tissues and fat on bovine tendon were first removed, then the bovine tendon was cut into 1 mm × 1 mm thin slices and washed utilising DI water. After that, the tendon was disinfected by immersion in 75% ethanol for 15 min. The tendon was then degreased in pure 2-propanol (1:15 w/v) on a shaker overnight. After thoroughly washing off 2-propanol utilising DI water, the tendon was soaked and continuously stirred in 0.5 M acetic acid (1:20 w/v) with 1% w/w pepsin for 48 h. After that, the resulting type I collagen-containing viscous solution was filtered using a 200 mesh strainer to remove undigested tendon. The filtered solution was adjusted to pH = 7.4 by 6 M NaOH, and the type I collagen fibrils self-assembled into larger fibres and hence gelation. The gel was centrifuged at 3,200 rpm for 15 min, and the supernatant was discarded. The gel-like sediment was collected and redissolved in 0.5 M acetic acid. After that, the solution was transferred to Amicon™ Ultra Centrifugal Filter Units with a 100 kDa cut-off filter and centrifuged at 3,500 rpm for 30 min to remove impurities and peptide debris. The resulting solution was then freeze-dried for 48 h to obtain lyophilised type I collagen powder.

### 2.2 Collagen yield and purity assessment

The dry content rate of the bovine tendon was first obtained via the weight discrepancy before and after dehydration. Triple replicates of 5.0 g wet tendon were dried in the oven at 60 °C for 24 h. Then, the dry tendon was weighed. The dry content rate (*R*_*dry*_) was calculated using the equation:

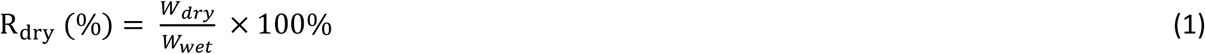

where

*W*_*dry*_ = dry specimen weight (g) after dehydration.

*W*_*wet*_ = wet specimen weight (g) before dehydration.

To obtain the yield of collagen extraction, three batches of 20.593 ± 0.462 g of wet tendon were utilised for extraction. Then, the total weight of lyophilised collagen powder was obtained, and the yield was calculated using the following equation:

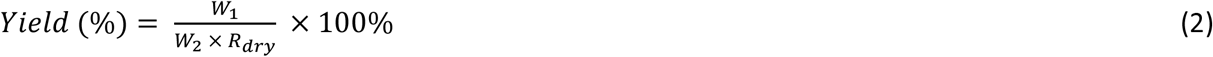

where

*W*_1_ = total weight (g) of lyophilised collagen powder.

*W*_2_ = weight (g) of wet tendon.

*R*_*dry*_ can be calculated from Equation (1).

To assess the purity of the extracted collagen from the above three batches, 0.1% w/v collagen samples in 0.5 M acetic acid from each batch (n = 3) was prepared for the assessments. The concentrations of total protein and total collagen in collagen samples were quantified using BCA total protein and total collagen assays, respectively, according to the manufacturer’s instructions. Three replicates were tested for each measurement. The purity was calculated using the equation:

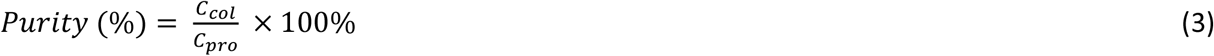

where

*C*_*col*_ = concentration (μg/mL) of total collagen in the collagen sample.

*C*_*pro*_ = concentration (μg/mL) of total protein in the collagen sample.

### 2.3 Methacrylation of type I collagen

To conduct methacrylation, type I collagen powder was dissolved in 0.5 M acetic acid to obtain a 0.6% w/v solution. Then, MAA was added at a volume ratio of MAA:collagen = 1:1000 with stirring at 500 rpm at room temperature. The mixture was stirred for 24 h, transferred to Amicon™ Ultra Centrifugal Filter Units with a 100 kDa cut-off filter and centrifuged at 3,500 rpm for 30 min to remove excessive MAA and byproducts. Then, the filtered solution was freeze-dried for 24 h to obtain lyophilized ColMA powder.

### 2.4 ColMA degree of substitution

To quantify the degree of substitution (DS), the 2,4,6-trinitrobenzene-sulfonic acid (TNBS) assay was conducted as previously described[12]. Briefly, 0, 8, 16, and 32 μg/mL glycine standard curves were plotted to obtain the primary amino group concentration. The extracted type I collagen and ColMA samples were separately dissolved at 100 μg/mL in 0.1 M sodium bicarbonate buffer. Then, 500 μL 0.01% TNBS in 0.1 M sodium bicarbonate buffer was added to 500 μL samples, followed by incubation at 37 °C for 2 h. After that, 250 μL of 1 M HCl and 500 μL of 10% w/v SDS were added to stop the reaction. The absorbance of the standard and test samples was measured at 335 nm on an Infinite M200 PRO plate reader (Tecan, Männedorf, Switzerland).

### 2.5 Fourier transform infrared spectroscopy and ultraviolet-visible spectroscopy

To obtain information on the functional groups of collagen samples, Fourier transform infrared (FTIR) spectroscopy and ultraviolet-visible (UV-Vis) spectroscopy were conducted. Lyophilized powder of commercial collagen, extracted collagen, and ColMA was utilised for FTIR spectroscopy on a Jasco FT/IR-4100 Spectrometer (Jasco UK Ltd, Essex, UK) using the micro KBr pellet method. The IR spectrum was scanned at room temperature from 4000 – 400 cm^-1^ with a resolution of 2 cm^−1^. To perform the UV-Vis scan, 10 μg/mL samples of commercial collagen, extracted collagen, and ColMA in 0.5 M acetic acid were separately prepared in a 1 mm thick quartzy cuvette. The UV-Vis spectra of collagen samples were recorded on a Jasco V-650 UV-Vis Spectrophotometer (Jasco UK Ltd, Essex, UK) in the range of 200 to 400 nm.

### 2.6 Western Blot

Lyophilized commercial bovine type I collagen (Sigma-Aldrich, Missouri, US), extracted type I collagen, ColMA, and commercial type B gelatin from bovine skin (Sigma-Aldrich, Missouri, US) were dissolved in 0.5 M acetic acid and stirred overnight at 4°C to obtain solutions with a concentration of 2 mg/mL. All the solutions were then mixed with NuPAGE™ LDS Sample Buffer (4X) at a volume ratio of 3:1 and reached a final concentration of 1.5 mg/mL. After incubation at 70°C for 10 min to denature proteins, 10 μL of each sample and 2 μL of the PageRuler™ Unstained Broad Range Protein Ladder were loaded into wells of the NuPAGE™ 10-well 3 – 8% Tris-Acetate Gel in the NuPAGE™ Tris-Acetate SDS Running Buffer. The electroblotting program was run in an ice bath and set at 70 V for 25 min followed by 100 V for 150 min. When finished, the gel was transferred to Coomassie Brilliant Blue staining solution (10% w/v Coomassie Brilliant Blue, 10% v/v absolute ethanol, and 3.5% v/v phosphoric acid in deionised water) and gently shaken on a plate shaker overnight at room temperature. After that, the stained gel was rinsed with deionised water on a plate shaker for 2 h, with fresh water changing at least three times, until the nonspecific staining background was almost washed off. The gel was then imaged utilising the ChemiDoc XRS + Gel Imaging System (Bio-Rad Laboratories, California, US).

### 2.7 Preparation of ColMA/PEGDA bioink and photoinhibitor dose adjustment

In a typical 10 mL ColMA/PEGDA bioink, 60 mg lyophilised ColMA powder was dissolved in 5 mL 0.5 M acetic acid and stirred overnight at 4°C to obtain a 12 mg/mL solution. After that, the ColMA solution was mixed with 75 μL of 40% w/v poly(ethylene glycol) diacrylate (PEGDA, Mn 700, Sigma-Aldrich, Missouri, USA). Fifty milligrams of LAP was dissolved in 1 mL of deionised water at 60°C for at least 15 min to allow thorough dissolution and then added to the mixture, followed by the addition of 0, 100, 250, 500, 750, or 1000 μL of Essential Natural Yellow Food Color (Curcumin E100, Waitrose, Acton, London). The volume was topped up to 10 mL by adding 0.5 M acetic acid and stirred for 10 min to obtain the homogeneous bioink. Appropriately spin the bioink to remove bubbles before printing. The 10 mm × 10 mm × 2 mm grid scaffolds were then printed for photoinhibitor dose optimisation. Printing was conducted on a LumenX 3D printer (Cellink, Gothenburg, Sweden) with a 405 nm wavelength laser, printed at 100 μm standard resolution on the z-axis, 20% laser intensity and 20 s exposure time for each layer.

### 2.8 Rheology of the ColMA/PEGDA bioink

All rheology experiments were performed on a Kinexus Prime pro+ rheometer (Netzsch, Selb, Germany) in parallel plate geometry (20 mm disk, 1 mm measuring distance) at room temperature. Oscillation frequency measurements were carried out at 0.5% shear strain and room temperature (25 °C) from 10 Hz to 0.1 Hz frequency. Single-frequency oscillation with changing temperature was carried out at 1% shear strain and 1 Hz frequency from 4 °C to 50 °C. For in situ UV curing, light from the Omnicure S2000 XLA Spot Cure Unit light source with 400 nm wavelength (Lumen Dynamix LDGI) was utilised to illuminate the underside of the bioink solution through the Kinexus UV Transparent Borosilicate Glass Plate. The UV intensity was set to 0.08 W/cm^2^, which is equivalent to a screen intensity of 20% on the LumenX 3D printer. The gelation kinetics were characterised by the evolution of the storage modulus (G’) and loss modulus (G’) over time. The UV light was turned on 60 s after recording and kept illuminating until the end of the 180 s recording. The gel point was defined as the crossover between G’ and G’’, indicating a transformation of the bioink from a liquid to solid state. Each test was repeated three times.

### 2.9 Swelling rate and degradation of the ColMA/PEGDA hydrogel

The water sorption capacity of the hydrogels was evaluated by immersing freeze-dried hydrogels with known weights in 1× PBS (pH = 7.4) at 37°C. Excessive liquid was wiped off using filter paper before weighing at predetermined time periods (20 min, 40 min, 60 min, 2 h, 4 h, 6 h, 12 h, 24 h, and 48 h). The swelling rate was calculated by using the following equation:

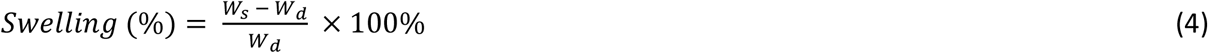

where

*W*_*s*_ = weight (mg) of hydrogel at the specific time point after immersion.

*W*_*d*_ = weight (mg) of freeze-dried hydrogel.

Similarly, hydrogels were fully soaked with 1× PBS (pH = 7.4) at 37°C for 24 h (day 0). Then, the hydrogels were weighed and incubated in PBS at 37°C for predetermined time periods (3, 7, 14, 21, and 28 days). Fresh PBS was replenished every week. The degradation of hydrogels was evaluated by the change in weight over time using the equation:

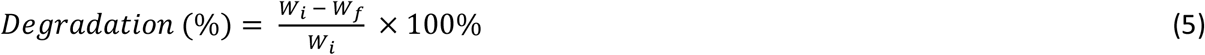

where

*W*_*i*_ = weight (mg) of hydrogel on day 0.

*W*_*f*_ = weight (mg) of hydrogel at later time points of measurement.

Three sample replicates were prepared for each time point of the above two evaluations.

### 2.10 Compression modulus of the ColMA/PEGDA hydrogel

The mechanical properties of DLP-printed ColMA/PEGDA scaffolds were characterised by unconfined compression tests using an Instron 5565 tester (Instron Ltd., Norwood, MA, US) with a 50 N load cell. Two types of ColMA/PEGDA cylinders (0.6% or 0.3% w/v ColMA + 0.3% w/v PEGDA, n=4, height = 5 mm, diameter = 10 mm) were fabricated utilising the LumenX 3D printer with a 405 nm curing wavelength, 100 μm standard resolution at the z-axis, 20% intensity, and 10 s exposure time for each layer, followed by 60 s post-curing using a 6 W, 405 nm visible light lamp. After that, the cylinders were immersed in PBS at 4 °C overnight, and excessive liquid on the surface was absorbed using filter papers before mechanical tests. For unconfined compression tests, each sample was placed between two compression plates and compressed at a displacement rate of 1 mm/min. The compression modulus was calculated as the slope of the linear region in the 0 – 10% strain range of the stress-strain curves, utilising the equation:

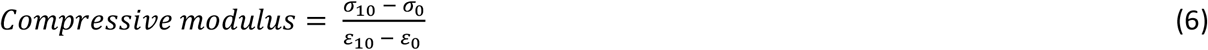

where

*σ*_10_ and *σ*_0_ = Compressive stress of the sample at 10% and 0% compressive strain, respectively.

*ε*_10_ and *ε*_0_ = 10% and 0% compressive strain, respectively, representing the change in height divided by the original height of the sample.

### 2.11 Printability of ColMA/PEGDA bioink and hydrogel microstructure

To test the printability of the ColMA/PEGDA bioink, a series of structures with fine details were fabricated. A necessary amount of ColMA/PEGDA bioink was loaded onto the petri dish of the LumenX 3D printer. Then, printing was performed with the same settings described in 2.7. Structures of the ear, microfluidic pads, and porous scaffolds were printed. The surface topography and porous structure of the printed hydrogels were investigated on a Zeiss Sigma 500 VP field-emission scanning electron microscope (FE-SEM, Zeiss, Germany). Samples were frozen at -80°C and then freeze dried, followed by gold coating before imaging at 2 kV in vacuum.

## 3 Results

### 3.1 Collagen solution yield and purity

The developed rapid preparation of collagen is shown in Figure 1. By utilising the method, the required time is shortened from 8 days to 4 days (Supplementary Figure 2), from the raw material to eventually obtained lyophilised powder. Ultracentrifugal filtration provides a more efficient way to separate unwanted small molecules and protein debris compared to conventional salting out plus dialysis. The extracted and purified collagen lyophilised powder exhibited a white to yellowish colour. The average dry content rate in the used bovine tendon was 37.8 ± 0.5% (SI Table 1). From 20.593 ± 0.462 g of wet tendon (containing 7.780 ± 0.078 g of dry content), 5.734 ± 0.021 g of lyophilised powder was obtained, resulting in a yield of 73.7 ± 0.9% (SI Table 2). In terms of purity, in the 1 mg/ml collagen solution sample, an average of 1017.95 ± 20.03 μg/mL total protein was measured, while 951.20 ± 27.57 μg/mL total collagen on average was confirmed. Hence, the developed rapid preparation of collagen shows the ability to reach a considerably high purity of 93.4 ± 2.7% (SI Table 3).

**Figure 1.**
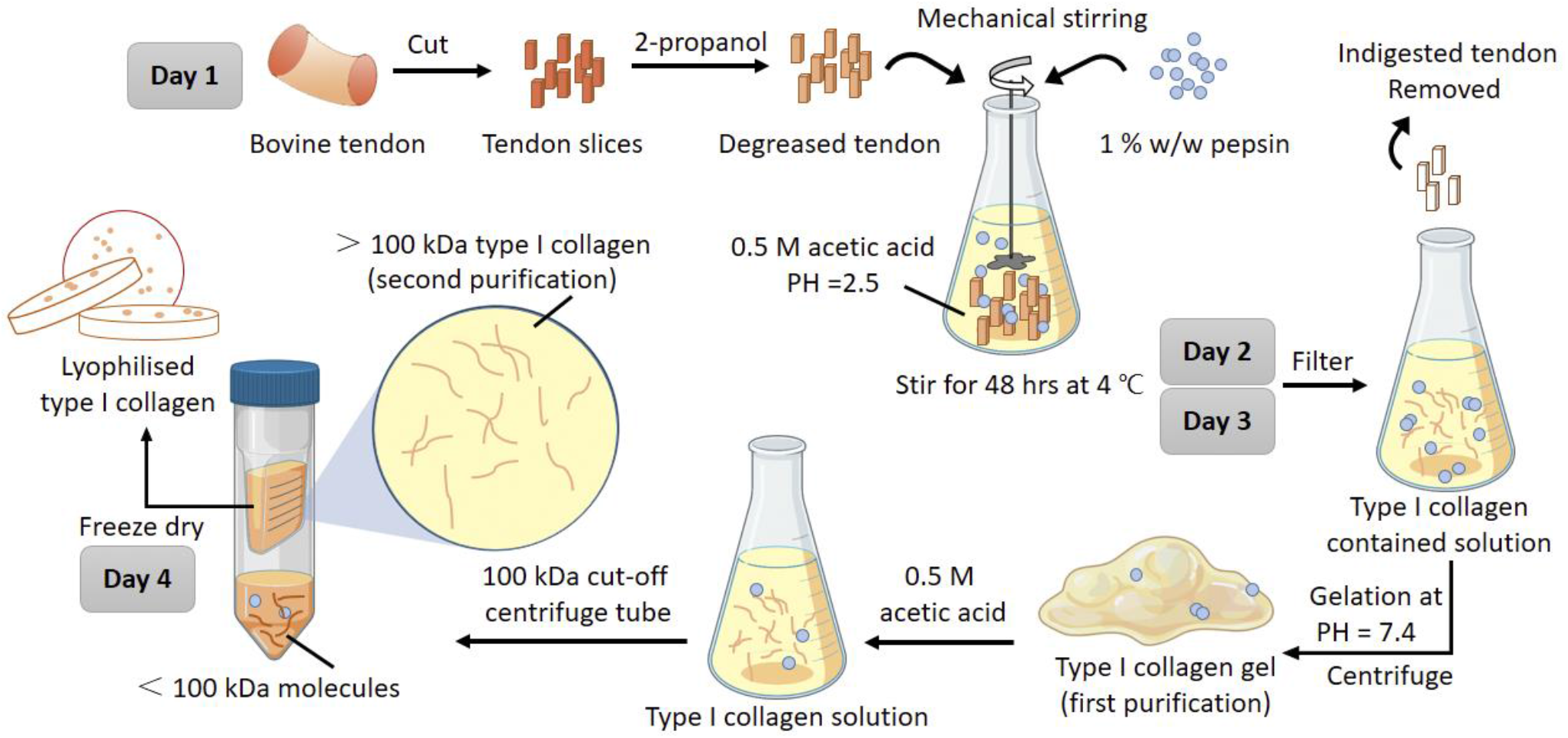
Schematic of the rapid preparation of collagen.

### 3.2 Methacrylation of type I collagen and degree of substitution

Methacrylation was carried out under acidic conditions (pH ≤ 2.5) at room temperature for 24 h. The MAA substitution reaction is illustrated in Figure 2a. The degree of substitution was evaluated utilising the TNBS assay (SI Table 4). The results showed that 100 μg/mL extracted type I collagen had equivalent amine groups of 2.88 ± 0.04 μg/mL glycine, from which 0.384 mmol free primary amino groups per gram collagen could be calculated. After methacrylation, ColMA was found to have equivalent amine groups of 0.34 ± 0.04 μg/mL glycine. Therefore, an average DS of 88.3 ± 1.4 % was reached. The crosslinking of ColMA can be triggered in the presence of the photoinitiator LAP and 405 nm visible light (Fig 2b).

**Figure 2.**
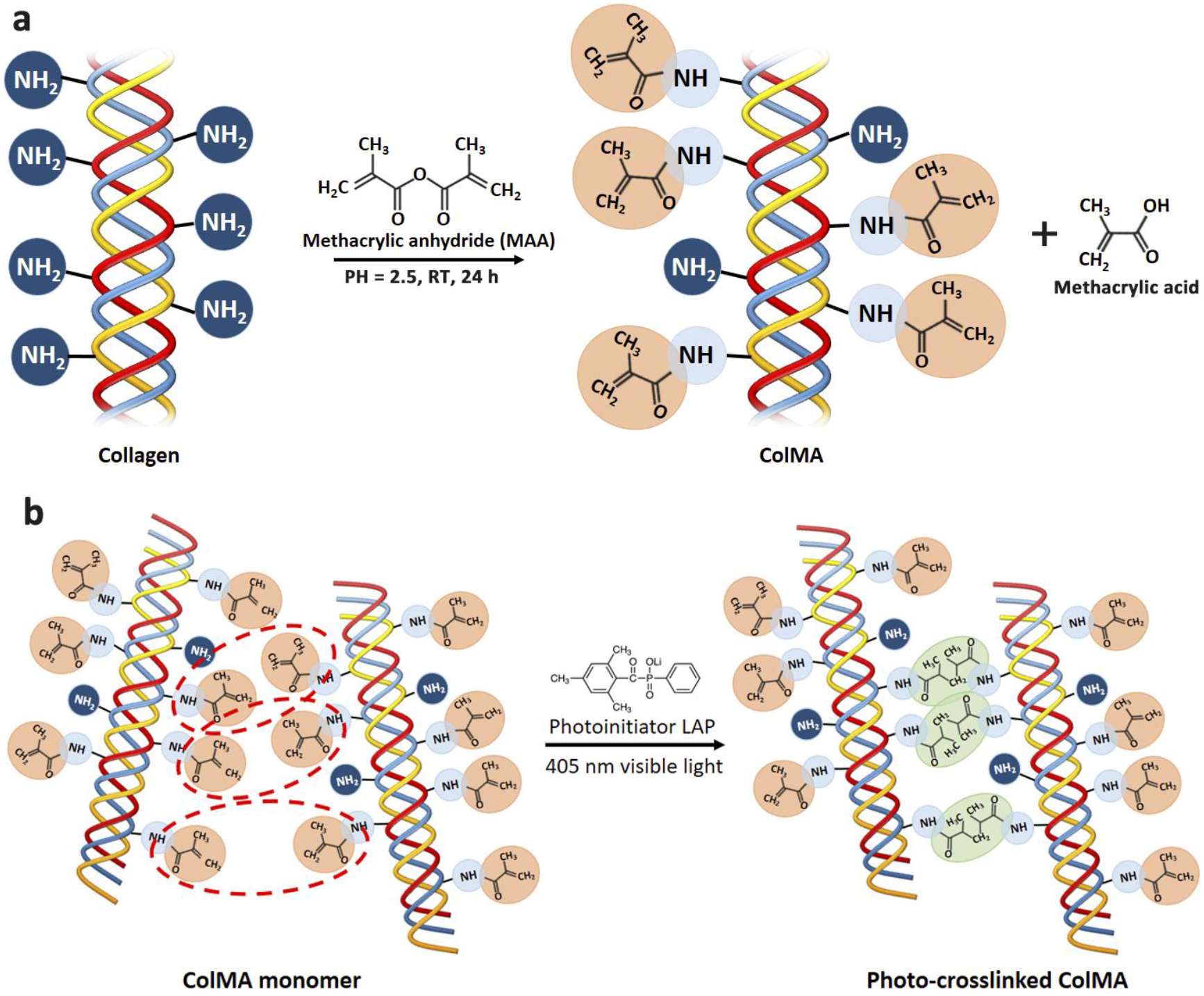
**(a)** The ColMA synthesis scheme: MAA reacts with the primary amines and leads to methacryloyl functionalization of collagen; **(b)** Schematic of the ColMA crosslinking reaction in the presence of LAP and 405 nm visible light.

### 3.3 FTIR and UV-Vis spectroscopy

Lyophilized powders of ColMA, extracted type I collagen, commercial type I collagen, and commercial type B gelatin were scanned by FTIR spectroscopy. The results (Fig. 3a) show that the extracted type I collagen, ColMA, and commercial type I collagen can be identified with characteristic peaks of amide A (3298 cm^-1^) and B (2922 cm^-1^) bands and amide I (1631 cm^-1^), II (1542 cm^-1^), and III (1236 cm^-1^) bands. Importantly, the peak of the amide III band indicates collagen with a complete triple helix structure. Commercial type B gelatin is the thermal degradation product of collagen without a triple helix structure; hence, the peak intensity of the amide III band is significantly reduced, and the peak of the amide A band is broadened. In UV-Vis (Fig. 3b), the extracted type I collagen, ColMA, and commercial type I collagen all show the characteristic peak of collagen at 230 nm, which can be attributed to the n→π* transitions of the C=O, -COOH, and CONH_2_ groups in the polypeptide chains of collagen[13]. The extracted type I collagen shows a broad peak at 250 nm – 280 nm, which indicates that it preserves more chromophore (primary amino group)[14] than the commercial type I collagen and is more efficient for the MAA substitution reaction. After MAA modification, the broad peak from 250 nm – 280 nm disappeared in the ColMA sample, indicating that most of the primary amino groups were substituted by MAA.

**Figure 3.**
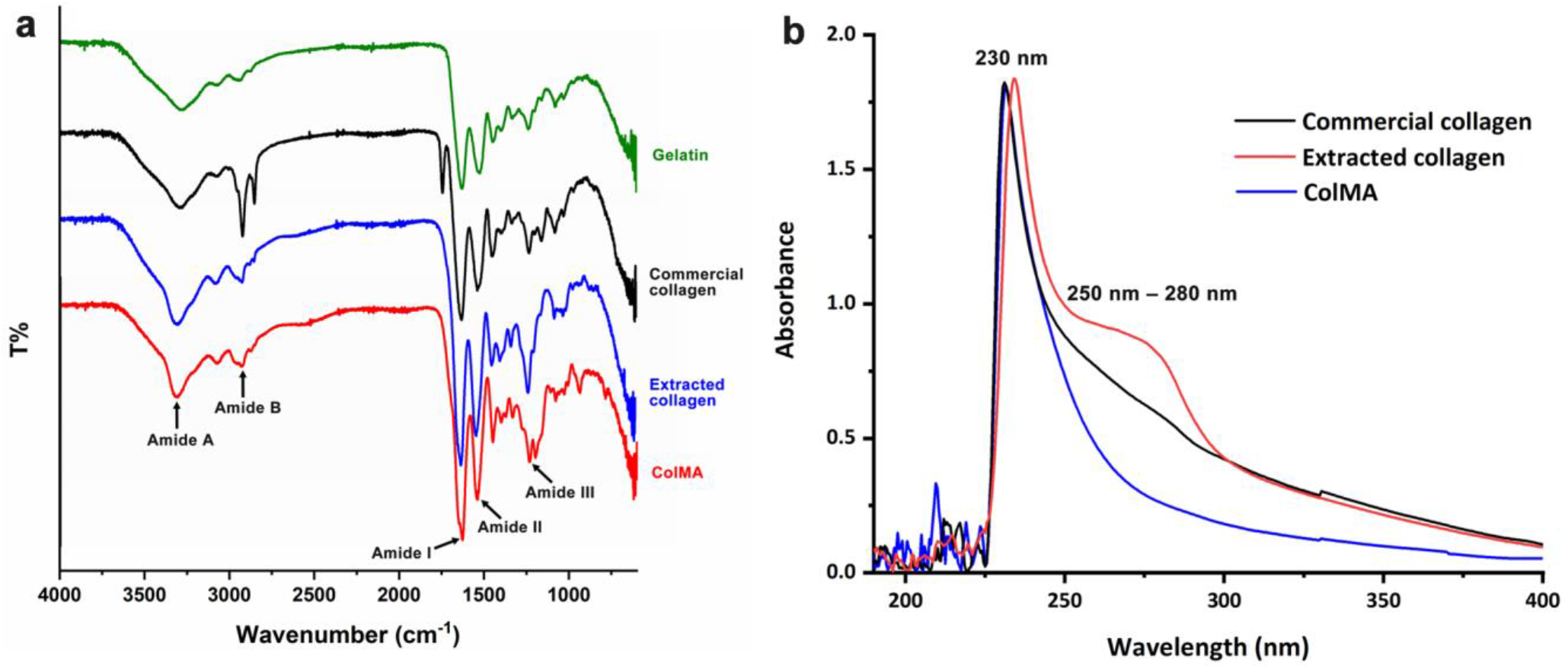
**(a)** FTIR and (**b)** UV-Vis spectra of different samples.

### 3.4 Western Blot

The Western blot electrophoresis results (Fig. 4. Uncropped gel see Supplementary Figure 3) showed that the bands of extracted type I collagen and ColMA were similar to those of commercial type I collagen, with identified α chain (130 kDa), β chain (α chain dimer, 250 kDa) and γ chain (α chain trimer, 400 kDa), and no other bands of small molecule proteins. In comparison, no bands over 100 kDa were found in the gelatin sample. The results further proved that the extracted collagen was high purity type I collagen, and the integrity of the triple helix structure was well preserved before and after MAA modification.

**Figure 4.**
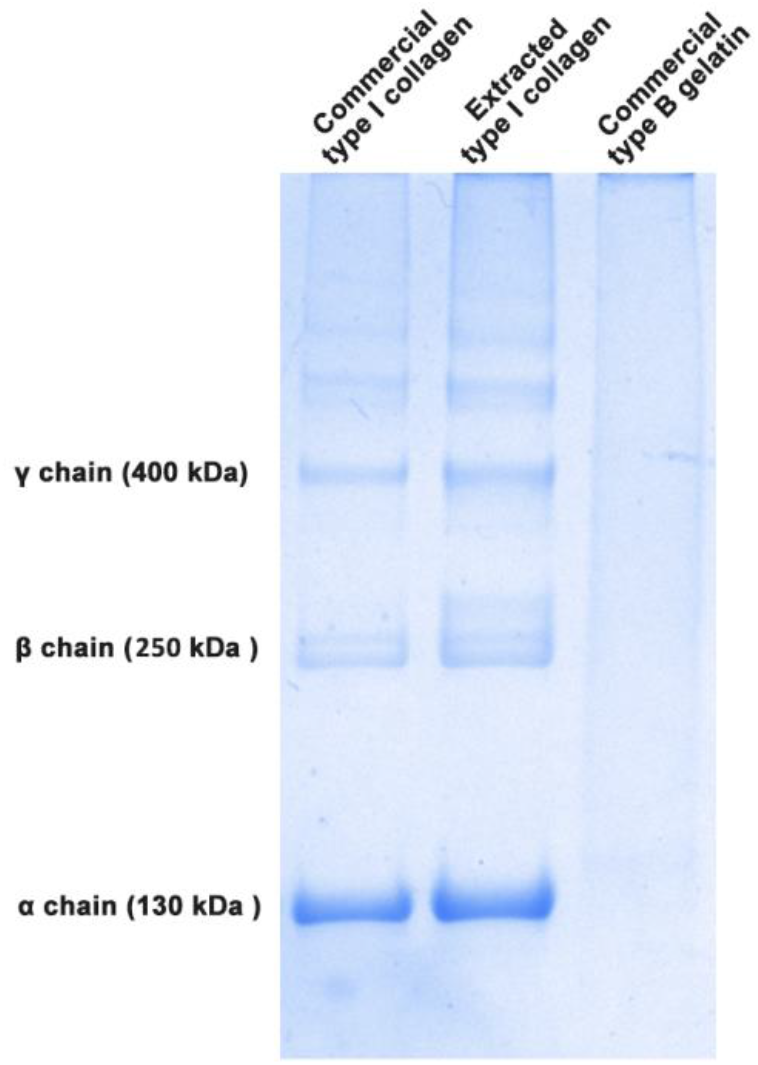
Western blot electrophoresis of different samples.

### 3.5 Preparation of ColMA/PEGDA bioink and photoinhibitor dose adjustment

The crosslinking reaction of ColMA in the presence of photoinitiator LAP and 405 nm visible light is depicted in Figure 4a. The addition of photoinhibitor is critical in the fine control of crosslinking depth. A lack of photoinhibitor results in crosslinking in unexpected areas, while excessive addition leads to insufficient crosslinking depth and thus incomplete structure or noncrosslinking. Blue light can be intensively absorbed by yellow objects. Therefore, a nontoxic yellow food dye is an ideal photoinhibitor. In Figure 5, by adding a gradient concentration of yellow dye, less than 5% v/v photoinhibitor caused over crosslinking within the grid, which significantly reduced the resolution of the structure. More than 5% v/v photoinhibitors lead to poor printability, as the missing part of the scaffolds displayed. The 5% photoinhibitor exhibits the best balance between excessive and insufficient crosslinking. It shows sharp structure and details while constraining the crosslinking just right on the projected image. Therefore, the optimised bioink formula of 10 mL ColMA/PEGDA bioink consists of 5 mL 1.2% w/v ColMA in 0.5 M acetic acid and 75 μL 40% w/v PEGDA (Mn 700). One milliliter of 5% w/v LAP, 500 μL of yellow food dye, and 3.425 mL of 0.5 M acetic acid.

**Figure 5.**
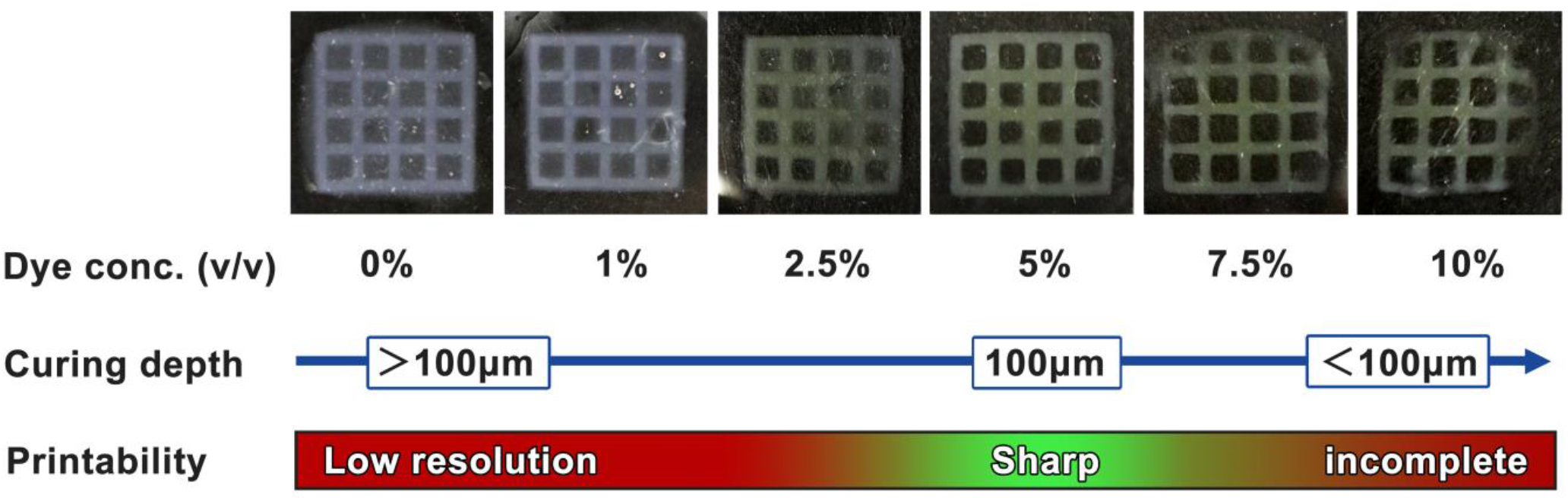
Printability of ColMA/PEGDA bioink with different additive quantities of photoinhibitor.

### 3.6 Rheology of the ColMA/PEGDA bioink

To understand the influence of temperature, visible light intensity, and exposure time on the ColMA/PEGDA bioink, the rheological profile and gelation kinetics of the bioink were obtained. The bioink displays a typical shear thinning behaviour at room temperature (Fig. 6a). Because the triple helix structure of collagen is thermally sensitive and its degradation is irreversible, a temperature sweep was carried out from 4°C to 50°C to investigate the thermal stability of the bioink. The results (Fig. 6b) show that the viscosity decreased when over 32°C, indicating the thermal stability for printing within the range of 4°C – 32°C. Normally, printing is conducted at room temperature (25°C), which is within the stable range of the collagen molecular structure. The gelation kinetic results (Fig. 6c) show that before curing, the samples were in a liquid phase. Upon exposure to 400 nm visible light, crosslinking was initiated, as demonstrated by the increasing storage modulus (G’). The loss modulus (G’’) intersected with the storage modulus (G’) after 10 s of curing (tan(δ) = 1), which defined the gel point of the bioink. It is shown that 10 s of exposure time under 0.08 W/cm^2^ is essential for bioink gelation. The curing nearly reached saturation when exposed for another 40 s. Therefore, a 10 s exposure time for each layer, followed by 60 s postcuring, is adopted in the following printing. The exposure intensity and time of postcuring exceeded the actual needs to ensure that full crosslinking could be achieved.

**Figure 6.**
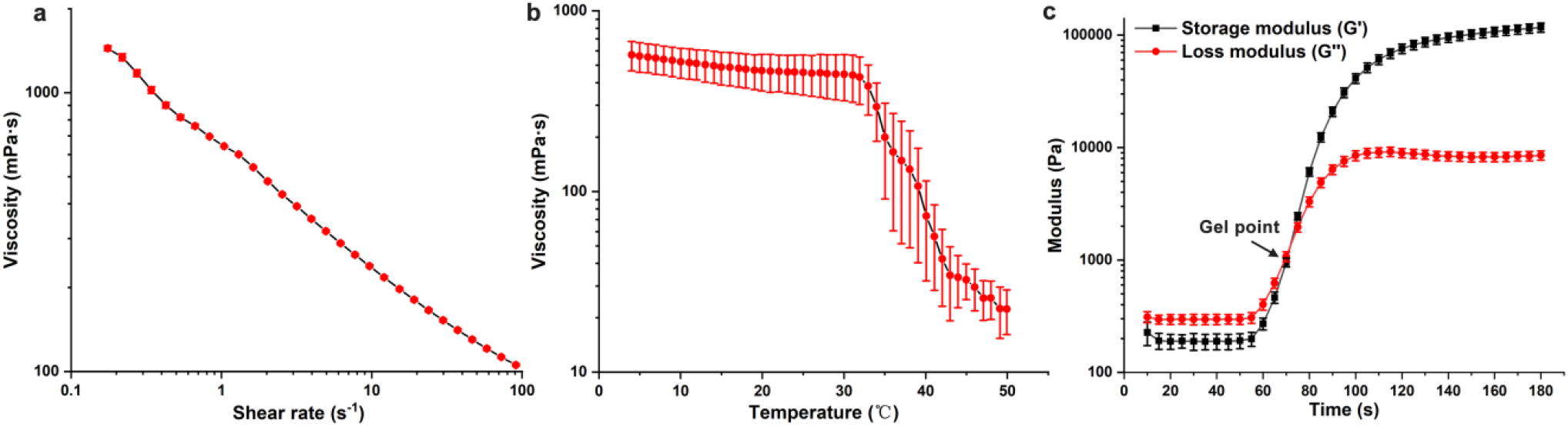
Rheology characterisation of the ColMA/PEGDA bioink (n = 3). Determination of **(a)** shear rate and viscosity; **(b)** temperature and viscosity; and **(c)** the evolution of the storage modulus (G’) and loss modulus (G’’) during in situ gelation. The arrow indicates the gel point, where G’ and G’’ crossover.

### 3.7 Swelling rate and degradation of the ColMA/PEGDA hydrogel

The swelling rate of hydrogels represents their water retention, gas exchange, nutrient transfer, cell encapsulation and ingrowth abilities[15, 16]. Figure 7a shows that the hydrogel intensively absorbed liquid in the first hour, then the sorption speed (slope of the curve) gradually slowed down in the following hours and nearly reached equilibration at 6 h. After that, the swelling rate steadily maintained over 800% and eventually reached 832.1 ± 69.2% at 48 h. Degradation is another important characteristic of hydrogel, indicating the essential resistance of degradation. The evaluation of degradation lasted for 28 days, which is considered to cover the length of most in vitro experiments. Figure 7b shows that the ColMA/PEGDA hydrogel degraded at a steady rate over time. A degradation rate of 35.6 ± 2.9% was found on day 28 of incubation in PBS.

**Figure 7.**
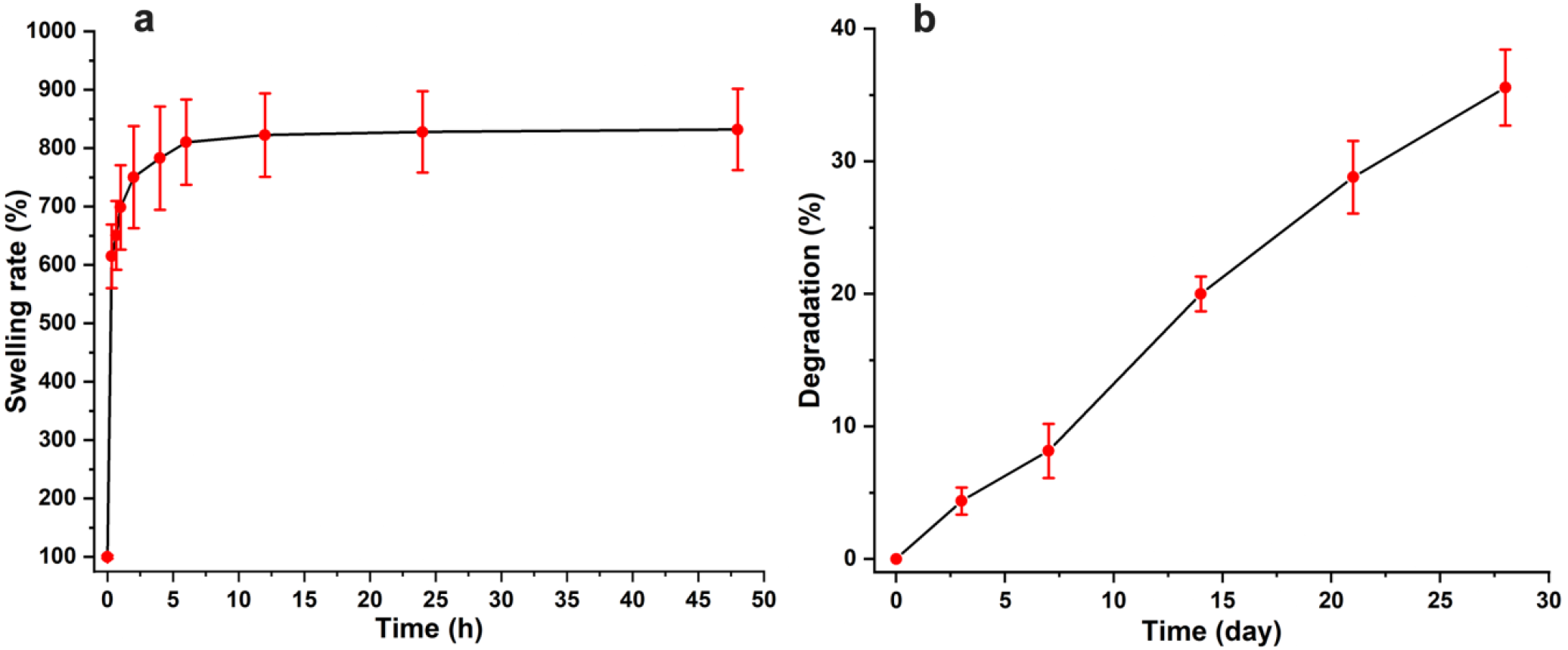
**(a)** Swelling rate of hydrogels in PBS over 48 h and **(b)** degradation of hydrogels in PBS over 28 days.

### 3.8 Compression modulus of the ColMA/PEGDA hydrogel

The compression modulus represents the stiffness of the printed ColMA/PEGDA hydrogel to withstand changes in length. The results (Fig. 8) show that the ColMA/PEGDA hydrogel reached 137.8 ± 19.0 kPa at low concentrations of 0.6% w/v ColMA and 0.3% w/v PEGDA. The collagen triple helix structure contributed remarkable mechanical strength to the crosslinking network. Moreover, the hydrogel showed tunable mechanical properties by adjusting the collagen additive quantity. Softer scaffolds with a compression modulus of 38.6 ± 10.6 kPa were obtained using the 0.3% w/v ColMA + 0.3% w/v PEGDA bioink. This illustrates the possibility that by precisely adjusting the concentration of the ColMA component, a broad range of mechanical properties can be achieved and are thus suitable for different purposes.

**Figure 8.**
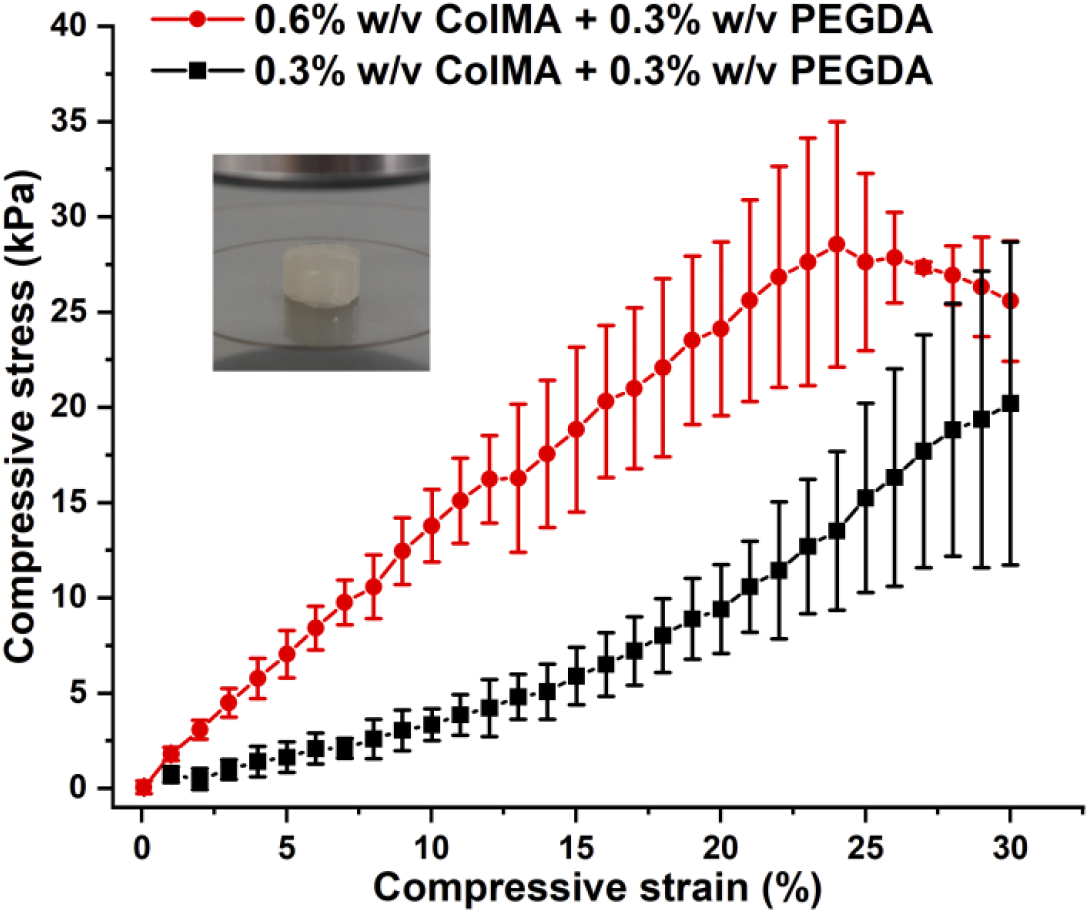
Compressive stress-strain profile of the ColMA/PEGDA hydrogels (n = 4) with different concentrations of the ColMA component. The illustration shows the 3D-printed hydrogel cylinder between the upper and lower plates of the testing instrument.

**Figure 9.**
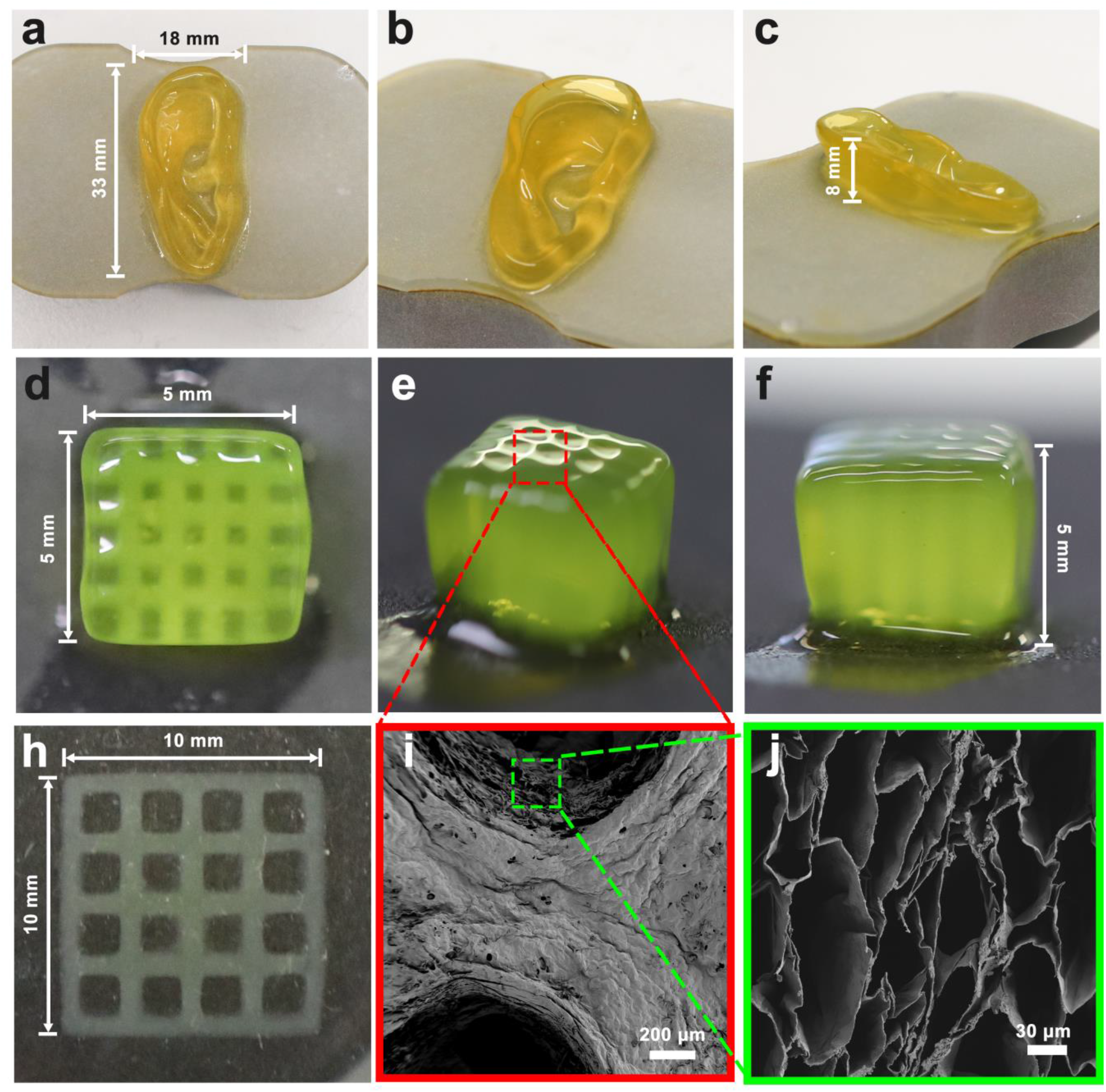
A series of 3D-printed ColMA/PEGDA hydrogels. **(a – c)** Ear large model; **(d – f)** Small cube with channels, showing a resolution of 500 μm at the x- and y-axes; **(h)** 10 × 10 mm medium-size scaffold (after postcuring, thickness = 1.5 mm) commonly used for 3D cell culture; SEM images of **(i)** surface topography and **(j)** porous structure at the cross section of the hydrogel.

### 3.9 Printability of the ColMA/PEGDA bioink

The printed ear and scaffolds all show sharp edges and fine details. There was no unexpected over-crosslinking or incomplete part of the structure. The ear model highlights the ability of large structure printing. It shows that even the complicated geometry is able to withstand its own weight, and no obvious deformation is witnessed while standing in the air. The small cube scaffold with channels exhibits the ability of high-resolution printing (up to 100 μm at the z-axis and 500 μm at the x- and y-axes in this study) and construction of a hollow structure utilising the developed bioink. The 10 × 10 mm scaffold is a popular model for in vitro experiments and thus used for the following biocompatibility tests. The SEM imaging at low magnification shows clear openings of the channel and rough topography of the hydrogel. The rough surface facilitates cell attachment and spreading[17]. The cross-sectional view at high magnification displays a porous structure with pore sizes of 50 – 150 μm. The porous structure and suitable pore size are critical to support cell migration and proliferation[18].

## 4 Discussion

DLP 3D printing is a promising technology, especially for sensitive bioinks, as curing can be performed at a relatively low temperature (4°C **–** 25°C), and the light for crosslinking is mild visible light. These conditions impose the minimum negative effects on the bioactive components of bioink. Nevertheless, DLP 3D printing requires bioink with low viscosity, as it significantly affects the driving abilities of the up-side-down printing platform. GelMA has long occupied the position of the most popular bioink for DLP. An indisputable reason is that gelatin achieves a balance among viscosity, thermal reversibility, and bioactivity. It can also work at a wide range of pH values, which brings effectiveness and convenience to the manufacturing process and cell mixing. Nevertheless, the poor mechanical properties of GelMA hydrogels are hard to ignore, which leads to a mismatched modulus with most tissues. From a biological perspective, collagen is a better option than gelatin. Type I collagen is a member of the collagen family. It accounts for one-third of the total protein in the human body and three-quarters of the skin weight[19] and is one of the most abundant protein components in the extracellular matrix[20]. Type I collagen consists of three polypeptide chains composed of a GLY-X-Y sequence. Among them, X and Y represent any amino acid, but mainly hydroxyticine and proline[19]. The intact triple helix structure of collagen endows it with outstanding mechanical properties and biological activity after crosslinking; thus, a wide scope of applications can be covered from soft brain tissue to hard bone tissue[21-23]. The integrity of the triple helix structure is the essential difference between collagen and gelatin. The poor mechanical performance and bioactivity of gelatin is largely due to degraded polypeptide chains[24]. Despite the biological advantages of collagen structure over gelatin, this special structure also sets hurdles in fabricating collagen-based bioink for DLP. To address most of the technical difficulties in collagen extraction, purification, bioink manufacturing, and DLP printing, we developed an optimised protocol covering the entire process, bringing solutions to collagen-based bioink for DLP printing.

The first step is collagen extraction. Type I collagen is abundant in sources and can be extracted from different types of animal tissues (bovine, porcine, chicken, marine, etc.)[25]. Various techniques can be used for extraction, usually including acid, alkali, enzymetic, or combined methods. In this protocol, the combined acid and enzymetic method was adopted. In most reported extraction methods, the extraction process involved enzyme digestion in acid, salting out, and dialysis, which took over one week to complete. The low-efficiency salting out and dialysis steps are the main reasons that make it time-consuming. To eliminate these steps, we utilised two-step purification. The first step is collagen gelation at neutral pH. This step takes advantage of collagen characteristics in that acid-soluble collagen fibrils can reversibly reassemble to form large fibres at physiological pH[26], and this phenomenon does not exist in noncollagen proteins. It allows a large proportion of crude collagen separation. After harvesting the gel and redissolving in acid, to more accurately remove unwanted components, collagen ultrafiltration units with a 100 kDa cut-off were utilised to separate large (type I collagen) and small molecules (enzymes, remaining noncollagen proteins, a part of liquid, and debris). This two-step purification achieved an average collagen purity of 93.4 ± 2.7% and a high yield (74.7 ± 0.9%) from three batches and can be finished within one hour. The FTIR and UV-Vis spectra show clear characteristic collagen-related peaks, and western blotting indicates no obvious impurity bands. Considering that the entire extraction process can be completed in 4 days, the purity and yield are even superior to those of previous reports extracting from the same source and using salting out and dialysis methods[27, 28], the developed method can be an optimised replacement for conventional methods. Additionally, the entire extraction process was performed at 4°C, which well preserved the bioactive triple helix structure, and more primary amino groups can take part in the MAA substitution reaction. Although the low temperature undeniably slowed pepsin hydrolysis, it is a trade-off for preserving the more important collagen molecular structure.

It has been widely reported that the primary amides on polypeptide chains of gelatin can be substituted by MAA to obtain GelMA[29]. The modified gelatin can be crosslinked by UV or visible light in the presence of a photoinitiator. As the source of gelatin with a similar molecular structure, the amide groups on the collagen peptide chains can theoretically also be substituted by MAA and photocrosslinked. To date, this idea has been proven feasible, and ColMA was produced[30, 31]. By regulating the collagen concentration, proportion of photoinitiator, UV intensity, and exposure time, the stiffness of the ColMA hydrogel can be controlled and made suitable for different applications. However, collagen-based bioink preparation is more complicated than gelatin-based bioink preparation, and the fabrication process has largely not been optimised, resulting in low efficiency and potential toxicity. The best reaction condition for MAA substitution is pH = 9 at 50°C for 1 **–** 3 h[12]. If the reaction conditions for collagen are acidic at 4°C, the efficiency will be significantly reduced. Considering that the acid condition is currently unable to be altered but needs to reach a high DS within an acceptable time span, we mildly raised the reaction temperature from 4 °C to 25 °C and appropriately prolonged the reaction time from the previously reported 3 h to 24 h. Under these conditions, a DS of 88.3 ± 1.4% was achieved at a significantly lower molar ratio (approximately 2.7 times molar excess) of the MAA over the primary amino group compared to the conventional GelMA synthesis, which usually uses a much higher excess amount of MAA (up to 44 times molar excess[32]) over the estimated amine groups. Much less MAA residuals and byproducts allow the ultrafiltration unit to remove most of the waste in 30 min instead of using conventional dialysis for days, which again significantly shortens the entire preparation time.

DLP printing cures the entire structure in a layer-by-layer manner. Therefore, the crosslinking depth is a critical parameter of the bioink. The only reported collagen-based bioink for DLP[33] is unable to control the crosslinking depth at each layer due to the absence of light inhibitors. Excessive crosslinking could occur exceeding the desired regions, resulting in blocked grids or channels. In this bioink formula, food-grade yellow dye was added as the photoinhibitor to restrain the curing depth with minimal cytotoxicity. It is found that 5% v/v addition ensures precise curing. The addition of PEGDA was reported to increase printing sharpness, and its giant PEG chain is able to form a noncovalent double network with the original hydrogel network[34]. The spontaneous penetration of the PEG network and its intrinsic biocompatibility make it a rational printing and mechanical tuner. The as-obtained ColMA/PEGDA bioink shows rheological features of shear thinning, stable viscosity below 32°C, and gel transformation after 10 s of exposure to visible light with an intensity of 0.08 W/cm^2^. The bioink displayed outstanding printability and flexibility in structure size from centimeters to hundreds of microns. The complicated ear sample was precisely printed without collapse or deformation while bearing its own weight in the air. The printed scaffolds reached a compressive modulus over 100 kPa after postcuring. Moreover, the hydrogel shows porous microstructure with pore size facilitating cell attachment, migration, and nutrition exchange[35]. The mechanical properties can also be easily adjusted by changing the additive proportion of ColMA. The tunable stiffness allows the developed ColMA/PEGDA bioink to be a promising candidate for a wide scope of biomedical applications.

The developed ColMA/PEGDA bioink has some drawbacks. The extracted collagen fibrils are still sensitive to pH. Self-assembly and gelation may be initiated when the pH increases. To keep free collagen fibrils and a stable viscosity, the bioink is maintained in acid (pH = 2.5), which prohibits cell premixing. This can be solved by rinsing the printed hydrogel in PBS to neutralise and then performing cell seeding. The concentration of collagen also has a great impact on the viscosity due to strong collagen fibril interactions[36], which hampers the use of more concentrated ColMA/PEGDA bioink for DLP printing. The bioink with 0.6 wt% ColMA is the highest concentration that we found that it can be printed without any difficulties. Any concentration above 0.6 wt% ColMA may result in higher viscosity and poorer printability (data not shown). This manuscript focuses on the fabrication and printing technique of the ColMA/PEGDA bioink. The biocompatibility test has not been shown in this manuscript but will be discussed in another specialised work in the future.

## 5 Conclusion

The rapid ColMA/PEGDA bioink preparation involves selective ultrafiltration, which greatly shortens the preparation cycle and is suitable for large-scale production. In methacrylation, a low MAA addition ratio ensures the lowest cytotoxicity from residuals and byproducts. The developed ColMA/PEGDA bioink retains its integrated bioactive triple helix structure and has low viscosity and good fluidity, which is suitable for DLP 3D printing. It reached high mechanical strength with a low concentration of collagen. The adjustable mechanical strength allows a wide range of medical applications. A series of structures were printed, showing great printability and the potential of individualised structures.

## Author Contributions

Conceptualisation, methodology, writing—original draft preparation, review, and editing, J.C. The author has read and agreed to the published version of the manuscript.

## Acknowledgements

The author thanks Prof. Wenhui Song for beneficial discussion, Chingyu Wang for consumable management, Lei Wu and Jinke Chang for equipment training and necessary assistance.

---

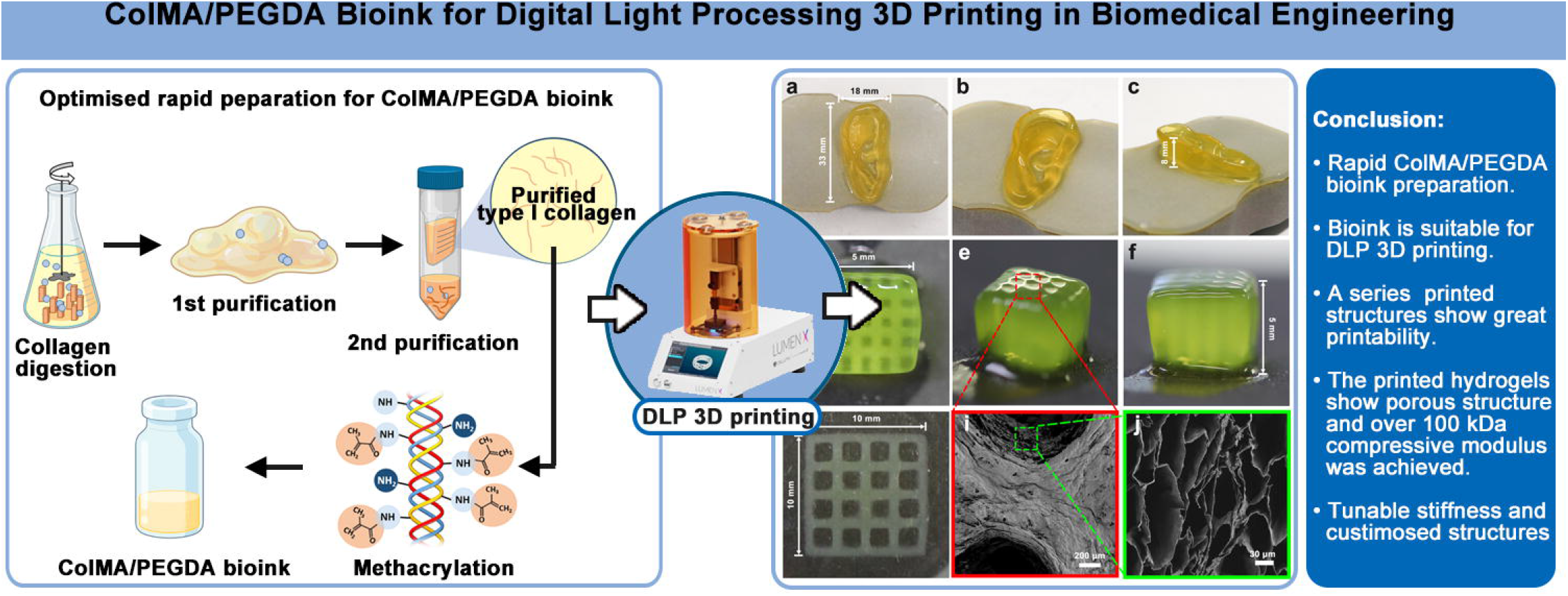

## References

[1] T.D. Ngo, A. Kashani, G. Imbalzano, K.T.Q. Nguyen, D. Hui, Additive manufacturing (3D printing): A review of materials, methods, applications and challenges, Composites Part B: Engineering 143 (2018) 172–196.

[2] K. Li, J. Rollins, E. Yan, Web of Science use in published research and review papers 1997–2017: a selective, dynamic, cross-domain, content-based analysis, Scientometrics 115(1) (2018) 1–20.

[3] J.K. Placone, A.J. Engler, Recent Advances in Extrusion-Based 3D Printing for Biomedical Applications, Adv Healthc Mater 7(8) (2018) e1701161.

[4] Y. Guo, H.S. Patanwala, B. Bognet, A.W.K. Ma, Inkjet and inkjet-based 3D printing: connecting fluid properties and printing performance, Rapid Prototyping Journal 23(3) (2017) 562–576.

[5] A. Sorkio, L. Koch, L. Koivusalo, A. Deiwick, S. Miettinen, B. Chichkov, H. Skottman, Human stem cell based corneal tissue mimicking structures using laser-assisted 3D bioprinting and functional bioinks, Biomaterials 171 (2018) 57–71.

[6] S. Naghieh, X. Chen, Printability-A key issue in extrusion-based bioprinting, J Pharm Anal 11(5) (2021) 564–579.

[7] W. Jeong, M.K. Kim, H.-W. Kang, Effect of detergent type on the performance of liver decellularized extracellular matrix-based bio-inks, 12 (2021) 2041731421997091.

[8] A. Persaud, A. Maus, L. Strait, D. Zhu, 3D Bioprinting with Live Cells, Engineered Regeneration 3(3) (2022) 292–309.

[9] A. Munaz, R.K. Vadivelu, J. St. John, M. Barton, H. Kamble, N.-T. Nguyen, Three-dimensional printing of biological matters, Journal of Science: Advanced Materials and Devices 1(1) (2016) 1–17.

[10] S.H. Kim, Y.K. Yeon, J.M. Lee, J.R. Chao, Y.J. Lee, Y.B. Seo, M.T. Sultan, O.J. Lee, J.S. Lee, S.I. Yoon, I.S. Hong, G. Khang, S.J. Lee, J.J. Yoo, C.H. Park, Precisely printable and biocompatible silk fibroin bioink for digital light processing 3D printing, Nat Commun 9(1) (2018) 1620.

[11] P.J.E.M. van der Linden, A.M. Popov, D. Pontoni, Accurate and rapid 3D printing of microfluidic devices using wavelength selection on a DLP printer, Lab on a Chip 20(22) (2020) 4128–4140.

[12] H. Shirahama, B.H. Lee, L.P. Tan, N.-J. Cho, Precise Tuning of Facile One-Pot Gelatin Methacryloyl (GelMA) Synthesis, Sci. Rep. 6(1) (2016) 31036.

[13] Z.-X. Xiang, J.-S. Gong, J.-H. Shi, C.-F. Liu, H. Li, C. Su, M. Jiang, Z.-H. Xu, J.-S. Shi, High-efficiency secretory expression and characterization of the recombinant type III human-like collagen in Pichia pastoris, Bioresources and Bioprocessing 9(1) (2022) 117.

[14] S. Prasad, I. Mandal, S. Singh, A. Paul, B. Mandal, R. Venkatramani, R. Swaminathan, Near UV-Visible electronic absorption originating from charged amino acids in a monomeric protein, Chem. Sci. 8(8) (2017) 5416–5433.

[15] N. Annabi, J.W. Nichol, X. Zhong, C. Ji, S. Koshy, A. Khademhosseini, F.J.T.E.P.B.R. Dehghani, Controlling the porosity and microarchitecture of hydrogels for tissue engineering, 16(4) (2010) 371–383.

[16] B. Maharjan, D. Kumar, G.P. Awasthi, D.P. Bhattarai, J.Y. Kim, C.H. Park, C.S. Kim, Synthesis and characterization of gold/silica hybrid nanoparticles incorporated gelatin methacrylate conductive hydrogels for H9C2 cardiac cell compatibility study, Composites Part B: Engineering 177 (2019) 107415.

[17] M. Ahearne, Introduction to cell-hydrogel mechanosensing, Interface Focus 4(2) (2014) 20130038.

[18] L.T. Vu, G. Jain, B.D. Veres, P. Rajagopalan, Cell migration on planar and three-dimensional matrices: a hydrogel-based perspective, Tissue Eng Part B Rev 21(1) (2015) 67–74.

[19] M.D. Shoulders, R.T. Raines, Collagen structure and stability, Annu. Rev. Biochem. 78 (2009) 929–58.

[20] C.M. Kielty, M.E. Grant, The Collagen Family: Structure, Assembly, and Organization in the Extracellular Matrix, Connective Tissue and Its Heritable Disorders 2002, pp. 159–221.

[21] A. Jain, M. Betancur, G.D. Patel, C.M. Valmikinathan, V.J. Mukhatyar, A. Vakharia, S.B. Pai, B. Brahma, T.J. MacDonald, R.V. Bellamkonda, Guiding intracortical brain tumour cells to an extracortical cytotoxic hydrogel using aligned polymeric nanofibres, Nature Materials 13(3) (2014) 308–316.

[22] A.M. Ferreira, P. Gentile, V. Chiono, G.J.A.b. Ciardelli, Collagen for bone tissue regeneration, 8(9) (2012) 3191–3200.

[23] C. Helary, I. Bataille, A. Abed, C. Illoul, A. Anglo, L. Louedec, D. Letourneur, A. Meddahi-Pellé, M.M. Giraud-Guille, Concentrated collagen hydrogels as dermal substitutes, Biomaterials 31(3) (2010) 481–490.

[24] S. Afewerki, A. Sheikhi, S. Kannan, S. Ahadian, A.J.B. Khademhosseini, t. medicine, Gelatin - polysaccharide composite scaffolds for 3D cell culture and tissue engineering: towards natural therapeutics, 4(1) (2019) 96–115.

[25] K.K.K. K. S. Silvipriya, A. R. Bhat, B. Dinesh Kumar, Anish John, Panayappan lakshmanan, Collagen: Animal Sources and Biomedical Application %J Journal of Applied Pharmaceutical Science, Issue: 3 2015.

[26] F.O. Schmitt, C.E. Hall, M.A. Jakus, Electron microscope investigations of the structure of collagen, 20(1) (1942) 11–33.

[27] H. Ju, X. Liu, G. Zhang, D. Liu, Y.J.M. Yang, Comparison of the structural characteristics of native collagen fibrils derived from bovine tendons using two different methods: modified acid-solubilized and pepsin-aided extraction, 13(2) (2020) 358.

[28] A. Sorushanova, I. Skoufos, A. Tzora, A.M. Mullen, D.I. Zeugolis, The influence of animal species, gender and tissue on the structural, biophysical, biochemical and biological properties of collagen sponges, J. Mater. Sci. Mater. Med. 32(1) (2021) 12.

[29] K. Yue, G. Trujillo-de Santiago, M.M. Alvarez, A. Tamayol, N. Annabi, A. Khademhosseini, Synthesis, properties, and biomedical applications of gelatin methacryloyl (GelMA) hydrogels, Biomaterials 73 (2015) 254–71.

[30] Q. Zhang, Q. Tang, Y. Yang, J. Yi, W. Wei, Y. Hong, X. Zhang, F. Zhou, X. Yao, H. Ouyang, Wound dressing gel with resisted bacterial penetration and enhanced re-epithelization for corneal epithelial-stromal regeneration, Applied Materials Today 24 (2021) 101119.

[31] P. Noohi, M.J. Abdekhodaie, M. Saadatmand, M.H. Nekoofar, P.M.H. Dummer, The development of a dental light curable PRFe-loaded hydrogel as a potential scaffold for pulp-dentine complex regeneration: An in vitro study, Int. Endod. J. (2022).

[32] B.H. Lee, H. Shirahama, N.-J. Cho, L.P. Tan, Efficient and controllable synthesis of highly substituted gelatin methacrylamide for mechanically stiff hydrogels, RSC Advances 5(128) (2015) 106094–106097.

[33] Z. Wu, J. Liu, J. Lin, L. Lu, J. Tian, L. Li, C. Zhou, Novel Digital Light Processing Printing Strategy Using a Collagen-Based Bioink with Prospective Cross-Linker Procyanidins, Biomacromolecules 23(1) (2022) 240–252.

[34] K. Chen, Y. Feng, Y. Zhang, L. Yu, X. Hao, F. Shao, Z. Dou, C. An, Z. Zhuang, Y. Luo, Y. Wang, J. Wu, P. Ji, T. Chen, H. Wang, Entanglement-Driven Adhesion, Self-Healing, and High Stretchability of Double-Network PEG-Based Hydrogels, ACS Applied Materials & Interfaces 11(40) (2019) 36458–36468.

[35] C.M. Murphy, M.G. Haugh, F.J. O’Brien, The effect of mean pore size on cell attachment, proliferation and migration in collagen–glycosaminoglycan scaffolds for bone tissue engineering, Biomaterials 31(3) (2010) 461–466.

[36] M. Shayegan, N.R. Forde, Microrheological Characterization of Collagen Systems: From Molecular Solutions to Fibrillar Gels, PLOS ONE 8(8) (2013) e70590.

