## Supplementary information for "ColMA/PEGDA Bioink for Digital Light Processing 3D Printing in Biomedical Engineering"

Jishizhan Chen^1,*^

*^1^ UCL Centre for Biomaterials in Surgical Reconstruction and Regeneration, Division of Surgery & Interventional Science, University College London, London, NW3 2PF, United Kingdom*

**Supplementary Information**

**^1^Table 1.** Wet and dry content in the bovine tendon samples.

|  | **Weight of wet tendon (g)** | | **Weight of dry tendon (g)** | | **Dry content rate (%)** |
| --- | --- | --- | --- | --- | --- |
| Sample 1 | 5.000 | 1.871 | | 37.4 | |
| Sample 2 | 5.000 | 1.922 | | 38.4 | |
| Sample 3 | 5.000 | 1.880 | | 37.6 | |
| Average ± SD | 5.000 ± 0.000 | 1.891 ± 0.022 | | 37.8 ± 0.4 | |

**Table 2.** Yield of collagen utilising the developed rapid method.

|  | **Weight of wet tendon (g)** | **Calculated weight of dry tendon (g)** | **Weight of lyophilised powder (g)** | **Yield (%)** |
| --- | --- | --- | --- | --- |
| Batch 1 | 20.134 | 7.751 | 5.711 | 73.68082 |
| Batch 2 | 21.225 | 7.887 | 5.73 | 72.6512 |
| Batch 3 | 20.421 | 7.703 | 5.761 | 74.78904 |
| Average ± SD | 20.593 ± 0.462 | 7.780 ± 0.078 | 5.734 ± 0.021 | 73.7 ± 0.9 |

**Table 3.** Purity of extracted collagen.

|  | **Theoretical conc. of sample (μg/mL)** | **Measured conc. of total protein (μg/mL)** | **Measured conc. of total collagen (μg/mL)** | **Purity (%)** |
| --- | --- | --- | --- | --- |
| Batch 1 | 1000.00 | 1038.567 | 951.202 | 93.4 |
| Batch 2 | 1000.00 | 990.824 | 917.431 | 90.1 |
| Batch 3 | 1000.00 | 1024.456 | 984.972 | 96.8 |
| Average ± SD | 1000.00 ± 0.00 | 1017.949 ± 20.027 | 951.202 ± 27.573 | 93.4 ± 2.7 |

**Table 4.** Equivalent glycine concentration in different samples.

|  | **Equivalent glycine conc. of collagen (μg/mL)** | **Equivalent glycine conc. of ColMA (μg/mL)** | **DS (%)** |
| --- | --- | --- | --- |
| Sample 1 | 2.83 | 0.30 | 89.7 |
| Sample 2 | 2.91 | 0.38 | 86.9 |
| Sample 3 | 2.90 | 0.34 | 88.3 |
| Average ± SD | 2.88 ± 0.04 | 0.34 ± 0.04 | 88.3 ± 1.4 |

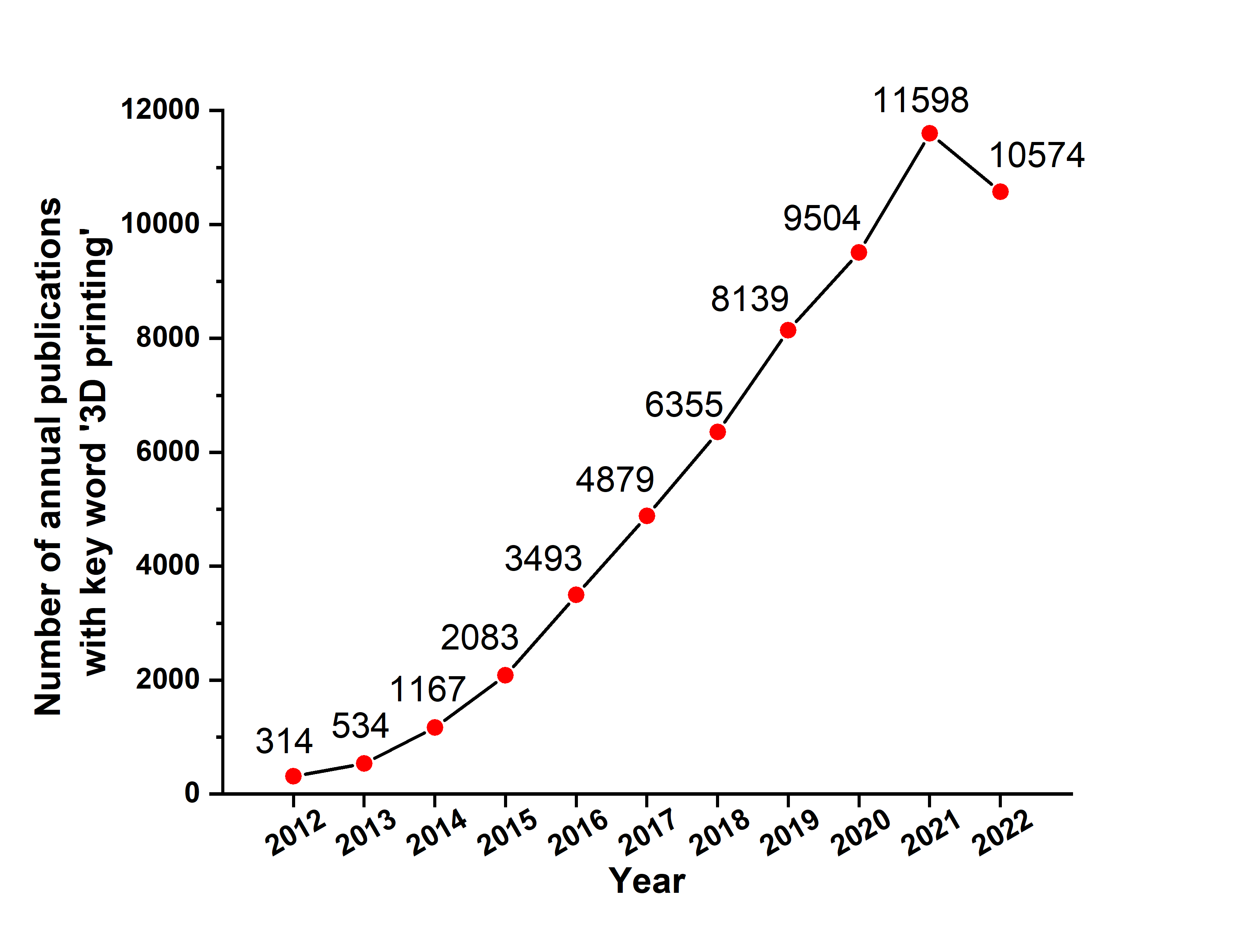

**Supplementary Figure 1.** Number of annual publications containing the key word ‘3D printing’, plotted based on the statistic data from the Web of Science, by 30th November, 2022.

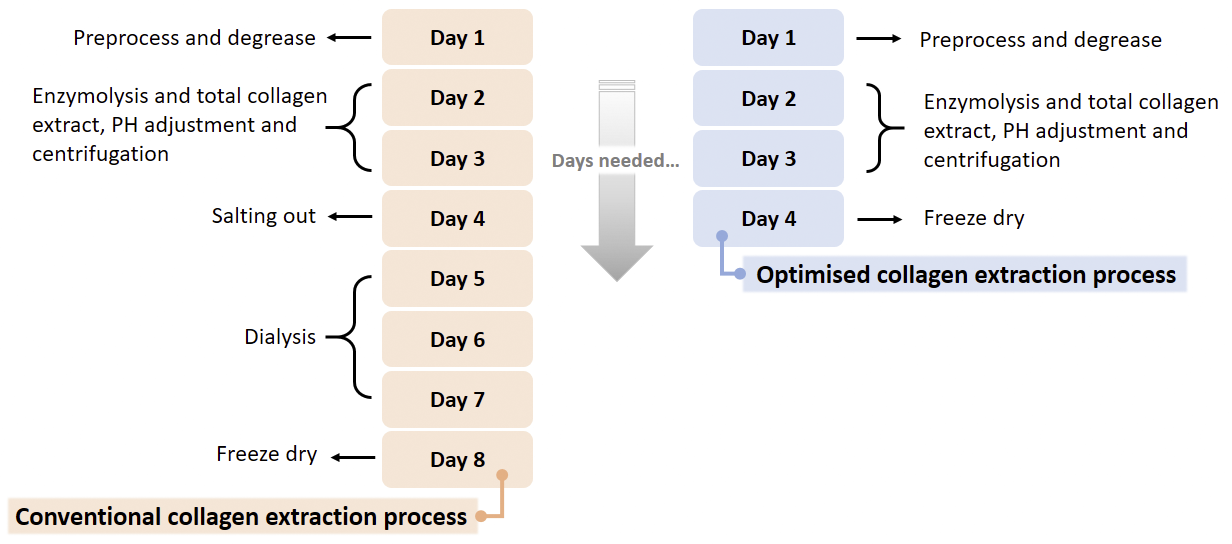

**Supplementary Figure 2.** Estimated elapsed time for the developed rapid collagen preparation and conventional method.

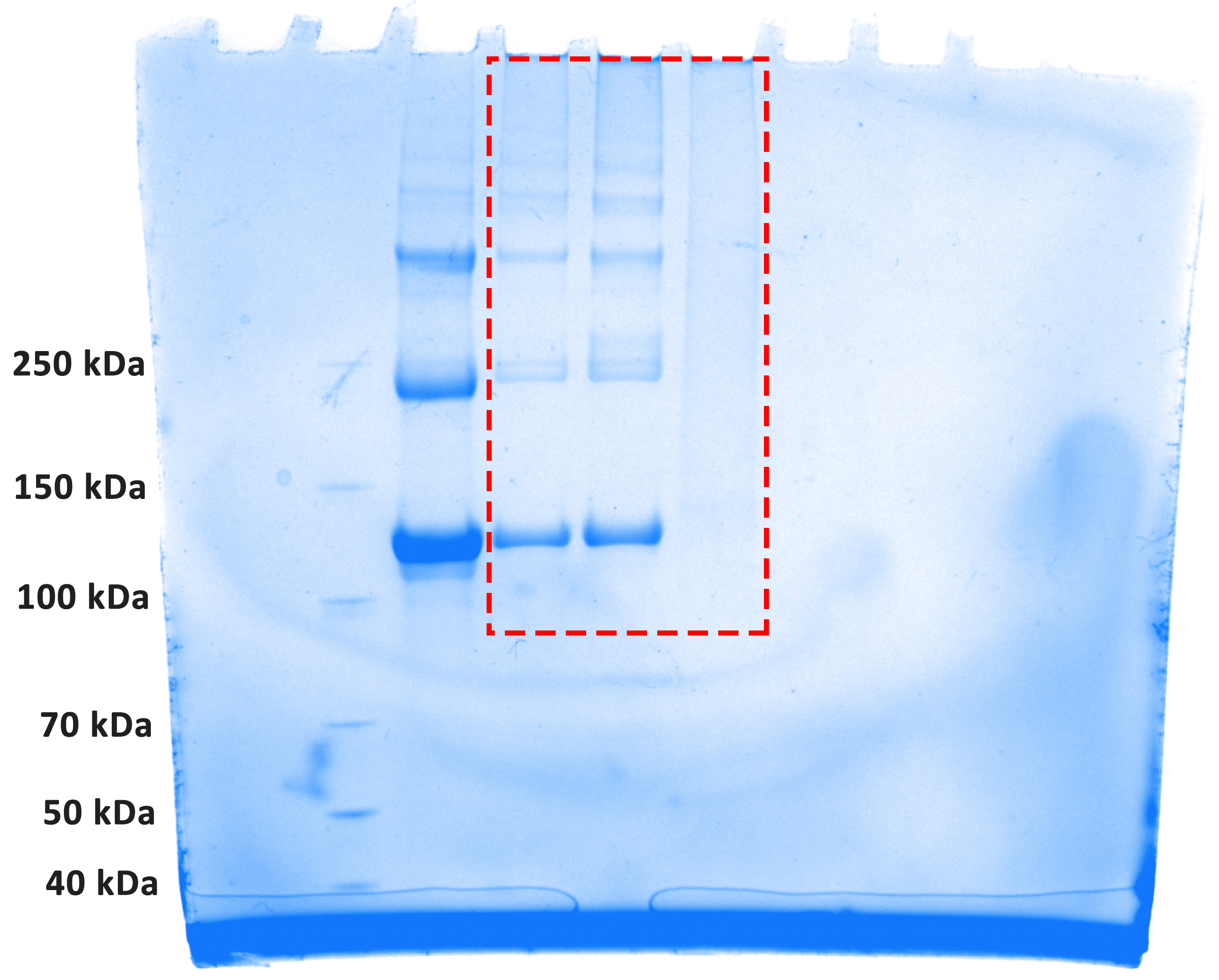

**Supplementary Figure 3.** Uncropped western blot electrophoresis gel. The red dashed box highlights the cropped area.

1	Sorushanova, A., Skoufos, I., Tzora, A., Mullen, A. M. & Zeugolis, D. I. The influence of animal species, gender and tissue on the structural, biophysical, biochemical and biological properties of collagen sponges. *J. Mater. Sci. Mater. Med.* **32**, 12, doi:10.1007/s10856-020-06485-4 (2021).
